# Alzheimer’s disease rewires gene coexpression networks coupling different brain regions

**DOI:** 10.1101/2022.05.22.492888

**Authors:** Sanga Mitra, Kailash B P, Srivatsan C R, Naga Venkata Saikumar, Philge Philip, Manikandan Narayanan

## Abstract

Connectome studies have revealed how neurodegenerative diseases like Alzheimer’s disease (AD) disrupt functional and structural connectivity among brain regions, but the molecular basis of such disruptions is less studied, with most genomic studies focusing on within-brain-region molecular analyses. We performed an inter-brain-region differential correlation (DC) analysis of postmortem human brain RNA-seq data available for four brain regions – frontal pole, superior temporal gyrus, parahippocampal gyrus, and inferior frontal gyrus – from Mount Sinai Brain Bank for hundreds of AD vs. control samples. For any two brain regions, our DC analysis identifies all pairs of genes across these regions whose coexpression/correlation strength in the AD group differs significantly from that in the Control group, after adjusting for cell type compositional effects to better capture cell-intrinsic changes. Such DC gene pairs provided information complementary to known differentially expressed genes in AD, and highlighted extensive rewiring of the network of cross-region coexpression-based couplings among genes. The most vulnerable region in AD, parahippocampal gyrus, showed the most rewiring in its coupling with other brain regions. Decomposing the DC network into bipartite (region-region) gene modules revealed enrichment for synaptic signaling and ion transport pathways in several modules, revealing the dominance of five genes (*BSN, CACNA1B, GRIN1, IQSEC2*, and *SYNGAP1*). AD cerebrospinal fluid biomarkers (AD-CSF), neurotransmitters, secretory proteins, ligand and receptors were found to be part of the DC network, suggesting how pathways comprising such signaling molecules could mediate region-region communication. A module enriched for AD GWAS (Genome-wide Association Studies) signals is also enriched for NF-κβ signaling pathway, a key mediator of brain inflammation in AD. Beyond modules, we also identified individual genes that act as hubs of AD dysregulation across regions, such as *ZKSCAN1* (Zinc Finger with KRAB And SCAN Domains) – this gene is known to be linked to AD in GWAS studies but via unknown mechanisms, and the specific DC interactions of *ZKSCAN1* found in this study can be used to dissect these mechanisms. Thus, our inter-region DC framework provides a valuable new perspective to comprehend AD aetiology.

## Introduction

The human brain connectome is comprised of a large-scale functional and structural network linking distinct brain regions, as evident from different neuroimaging techniques^1,2^. Brain functions in health rely on the connectome map, and disease can lead to its rewiring and disruption^3^. Structural connectome influences functional connectome shaping brain region specific activity^4^. Investigating the brain connectome has helped understand the abnormalies in brain connectivity of progressive neurodegenerative diseases such as Alzheimer’s disease (AD)^5,6^, which is characterized by the extracellular amyloid beta (Aβ) plaque development and intracellular neurofibrillary tangles (NFTs) formation at the molecular level, and manifests as memory loss, cognitive dysfunction, behavioural abnormalities, and social disorders^7^ at the clinical level. Genome-wide association studies (GWAS) of clinically diagnosed AD have been performed to identify risk loci and potential causative genes^8,9^, but which brain regions and mechanisms these genes act through is not fully characterized, and hampers therapeutic interventions aimed at slowing down or halting neuronal loss associated with AD.

Genomic studies are becoming instrumental to understand the molecular basis of neural circuits^10,11^ connecting different brain regions in health and disease. Brain functional connectivity is known to be under genetic control^12,13^. Recent studies are trying to link gene expression to connectome data^14,15,16^. Various studies have employed genome-wide gene expression (transcriptomic) analysis to elucidate gene regulatory interactions operating within brain tissue or regions of healthy/diseased individuals^7^. The effect of AD on different cortical regions has also been studied using gene expression-based molecular network analysis^17^. There are also several established differential expression (DE) studies, which address disease-induced change of expression level of individual genes in a region-specific manner. But these and other current AD gene expression studies^18,19^ have mainly focused on within-tissue/within-region analysis to provide insights into disease genes/processes. Therefore, the molecular mechanisms supporting inter-brain-region functional and structural connectivity, i.e., gene-expression coordination across brain regions, especially in neurodegenerative disease states, remain undefined.

To understand the gene-gene couplings across brain regions under normal conditions and their change in disease conditions, we construct a differential correlation (DC) network across different brain regions. We account for cellular composition effects in the data to better capture cell intrinsic changes in disease. The DC network is comprised of gene pairs whose correlation strength is altered (lost or gained) in disease (AD) group compared to control group. Interestingly we noted that each brain region uses a unique set of genes while interacting with genes of other brain regions. The DC gene pair rewiring is most prominent for coupling of one of the regions, parahippocampal gyrus, with other brain regions, in accordance with earlier studies on vulnerability^20^ or white matter degeneration^21^ of different brain regions.

By assessing how AD rewires this network of inter-region correlations (gene-gene coexpression patterns) through bipartite network analysis, we have uncovered novel interactions and identified synaptic signaling and ion transport pathways as the most affected biological processes across brain regions. This pair of pathways is found to be enriched in two DC modules, one module located in frontal pole and the other being in parahippocampal gyrus, revealing the dominance of five genes (*BSN, CACNA1B, GRIN1, IQSEC2*, and *SYNGAP1*). Systematically screening the DC network for hub genes revealed an AD-GWAS signal enriched gene *ZKSCAN1*^22^ as a dominant connector, indicating its probable role and mechanism in AD pathogenesis. In order to formulate mechanistic hypotheses underlying these AD-induced DC relations, we performed customized over representation analysis for the bipartite modules using molecular components essential for communication between proximal/distal cells or regions, such as ligand-receptor molecules, AD-CSF markers (proteins that are known to be biomarkers for AD), secreted proteins (secretome), and neurotransmitters-neuroreceptors (neurotransmission). We detected these different types of signaling molecules, specifically neurotransmitter, in our DC network, denoting the plausible mechanism of information transfer across brain regions. Further preliminary analyses indicate validation of our DC relations in an independent AD cohort. Taken together, our results furnish with vital molecular information about AD pathogenesis that may help in promoting AD therapeutics.

## Results

### Gene pairs between brain regions are rewired in AD pathology

Study of the rewiring of the coexpression network observed in the diseased group compared with the healthy control group is a great starting point for understanding the gene level disruption caused by the disease condition^23^. The differential correlation analysis identifies such gene pairs that have either significantly gained or lost correlation in disease compared to the control group. Using RNA-seq data from Mount Sinai Brain Bank (MSBB) of four brain regions, namely Brodmann Area (BM10) - frontal pole (FP), BM22 - superior temporal gyrus (STG), BM36 - parahippocampal gyrus (PHG), and BM44 - inferior frontal gyrus (IFG); and inter-region DC analysis (**Fig. 1a**), we noted significant rewiring of gene pairs between two brain regions in AD compared to CTL samples (**Table 1, Fig. 1b**).

**Table 1.** For each inter-brain-region comparison, number of DC gene pairs (edges) and unique DC genes (nodes) in the DC network detected at FDR 1% are reported (A DC gene is any gene participating in a DC relation; BR1 and BR2 stands for Brain Region 1 and 2 respectively; and DC Dysregulation index is the percentage of detected DC pairs out of all tested gene pairs).

| **Brain Region Pairs (BR1-BR2)** | **Total DC edges** | **BR1- # DC Genes** | **BR2- # DC Genes** | **DC Dysregulation index** |
| --- | --- | --- | --- | --- |
| FP-STG | 2961 | 2013 | 1844 | 2.54 |
| FP-PHG | 2629 | 1597 | 1409 | 5.41 |
| FP-IFG | 9962 | 4609 | 4235 | 3.40 |
| STG-PHG | 6274 | 2580 | 2549 | 8.94 |
| STG-IFG | 8179 | 3297 | 3548 | 5.24 |
| PHG-IFG | 12979 | 3642 | 4202 | 14.01 |

**Figure 1.**
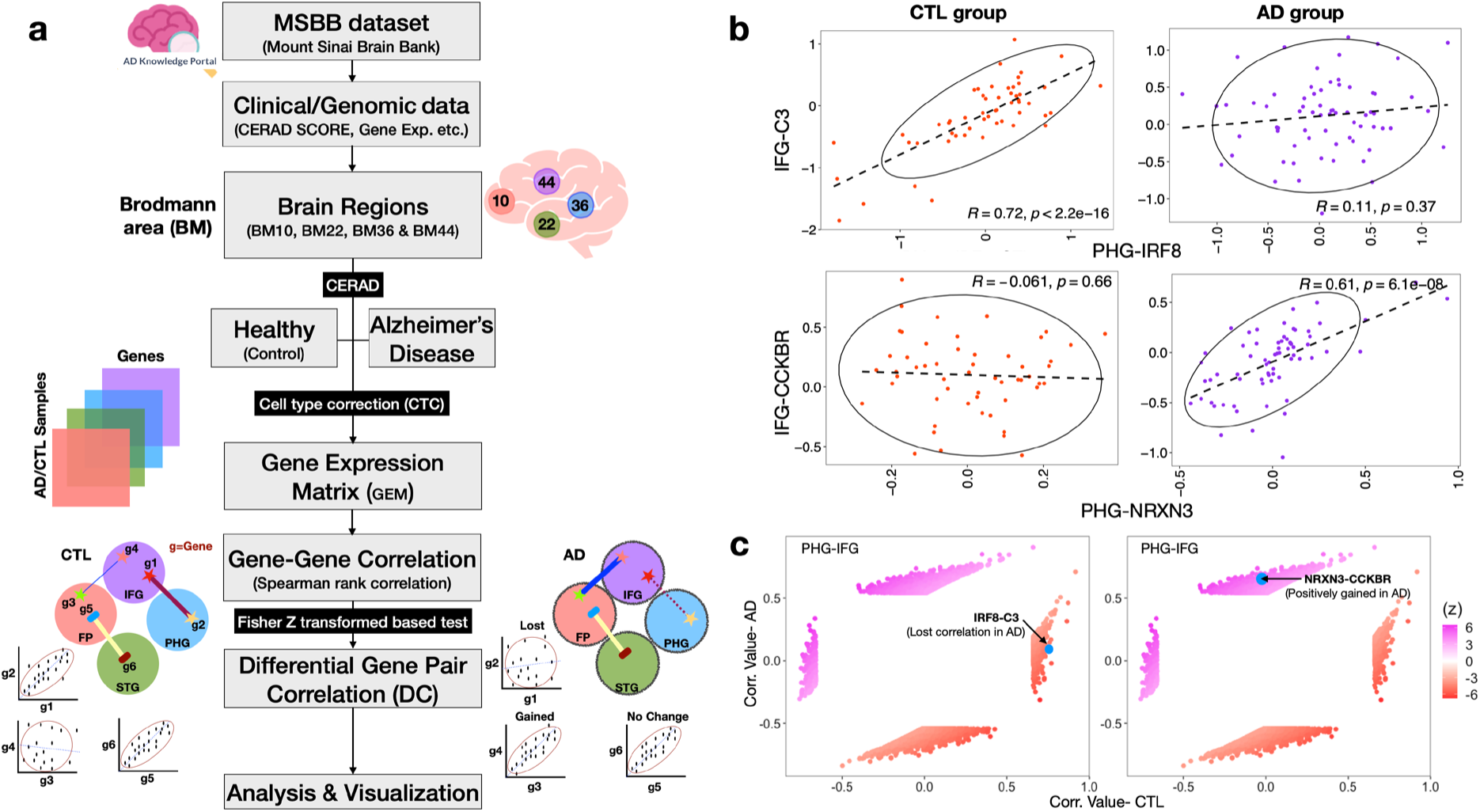
Gene pairs are differentially correlated (DC) in brain inter-region comparison in AD pathology. a. Schematic summary of the methodology - To understand inter-brain region dysregulation we obtained bulk tissue gene-expression datasets from MSBB for four different brain regions (for details see text); grouped them into AD (Alzheimer Disease) and healthy (control, CTL) samples based on CERAD score, and computed gene-gene Spearman’s correlation between every pair of brain regions separately in the AD group and CTL group. By correcting the expression data for cell-type composition effects (cellular deconvolution) before DC analysis, the confounding influence of cell-type proportions causing DC patterns is mitigated. b. We note that gene pairs are rewired between AD and CTL, leading to altered gene networks in AD pathogenesis. Two categories of changes, a gain of correlation (*IRF8* - *C3*) and loss of correlation (*NRXN3*-*CCKBR*) were detected. c. The DC pairs form four distinct clusters in a scatter plot representing the different categories of changes detected from DC analysis between AD and CTL. Comparing the size of the two gained correlation clusters (both positively and negatively gained) with the lost correlation clusters, there are more gained correlation gene pairs (here, absolute correlation cut-off of at least 0.4 is used to call a gene pair correlated).

Interestingly, our inter-region comparison of PHG-IFG, the two most vulnerable of these four regions according to an earlier study^20^, shows the maximum rewiring of gene pairs (12,979) compared to the other five inter-region comparisons. Additionally, the DC Dysregulation index, which captures the prevalence of DC pairs, attained a maximum of 14.01 for the PHG-IFG region pair, and minimum of 2.54 for FP-STG. This indicates that gene interactions between the two most vulnerable brain regions in AD are most affected, whereas those between less vulnerable regions (FP-STG) are the least affected (**Table 1**). Notably, the DC dysregulation index also declined depending on the decreasing vulnerability rank of the brain regions interacting with PHG. Note that PHG was reported as the most vulnerable site in AD^20^, and also exhibited prominent white matter tract degeneration^21^.

For each significant DC gene pair, we observed that the gene pair has either lost or gained correlation in AD compared to CTL (**Fig. 1b**). Further, delineating a DC pair based on a z-score threshold yielded 4 categories (**Fig. 1c, Suppl. Fig. 1b & Suppl. Table 1**), which can be grouped into 3 classes: gained positive correlation, lost correlation, and gained negative correlation. The position and the class of DC for the gene pairs, IRF8-C3 and *NRXN3-CCKBR* from PHG & IFG, are highlighted in Fig 1c. *NRXN3* and *C3* are already reported as AD-CSF (Cerebrospinal Fluid) biomarkers. This indicates that CSF biomarkers, which are known to help in brain communications along with ISF (Interstitial Fluid), may help in rewiring gene pairs in AD^24^. Further, assessing the overlap of DC gene sets from all six inter-region comparisons, we realized only a few gene pairs are common across these comparisons (**Suppl. Fig. 1d**). This indicates that the rewiring of gene pairs varies based on which pair of brain regions are taken under consideration for DC analysis.

### DE genes do not drive DC gene pairing

Since DE-based vulnerability index from an earlier study and our DC-based dysregulation index provide similar rankings for how the different brain regions are affected by AD, we wanted to check if DC results are driven by DE or if they complement DE. We checked the overlap between DC and DE genes for every inter-region comparison. In this study, we have used cell type corrected (CTC) DEGs for comparison since CTC data is also used for computing DC. Significantly altered CTC-DEGs are compared with DC genes participating in all six inter-region comparisons at FDR 0.05, 0.1, and 0.2. Even at a relaxed cut-off of FDR 0.2, more than 90% of DC gene pairs are not driven by DEG **(Table 2)**. While testing how many DC edges overlap with DEGs, it became evident that 9% of edges are driven by DEG for FP-STG comparison, whereas for PHG-IFG, only 1% are affected. This ensures that DEGs do not confound DC relations.

**Table 2.** DC (FDR 1%) vs. DEG (FDR 20%) overlap. -A DC edge is said to be driven by DE if any of the two genes in this DC edge or both genes are DE.

| Brain Region Pairs | Total DC edges | DC edges not driven by DE | DC edges driven by DE |
| --- | --- | --- | --- |
| FP-STG | 2961 | 2695 (91%) | 266 (9%) |
| FP-PHG | 2629 | 2460 (94%) | 169 (6%) |
| FP-IFG | 9962 | 9221 (93%) | 741 (7%) |
| STG-PHG | 6274 | 6056 (97%) | 218 (3%) |
| STG-IFG | 8179 | 8006 (98%) | 173 (2%) |
| PHG-IFG | 12979 | 12838 (99%) | 141 (1%) |

### FP-centred set of DC gene pairs replicates in an independent cohort

After determining the significant rewiring of gene pairs in inter-brain regions from MSBB cohort, we wanted to validate the DC patterns in another independent cohort to gain confidence in our findings. We tested for replication using another cohort data from Harvard Brain Tissue Resource Center (HBTRC), where different regions have been profiled^25^; specifically two cortical regions, visual cortex (VC, BM17) and dorsolateral prefrontal cortex (DLPFC, BM9), profiled across 300 samples from HBTRC (116 CTL and 184 AD) were used for replication. Only FP region was common between the two cohorts, i.e., DLPFC in HBTRC was closet to FP in MSBB. So, we focussed on FP-centred DC pairs and tested only them for replication in the HBTRC cohort. We followed this strategy as we couldn’t find another human cohort with transcriptomic data on the same brain regions as MSBB at sufficient sample sizes (>= 30). Another challenge we faced during replication is that DC gene pairs in MSBB cohort across brain inter-region comparison share almost no similarity at FDR 1% (**Suppl. Fig. 1d**). So, we decided to relax the cut-off from 1% to 30% to obtain sufficient number of common DC pairs between FP-STG and FP-PHG (see detailed schematic of replication in **Suppl. Fig. 2**). This resulting set of observed FP-centered DC edges (195,102 edges at FDR 30%) had an overlap of 407 pairs against HBTRC DC pairs (105,249 DC pairs at FDR 1%), and this replication overlap was significantly better than that of random gene pairs with the same size and network structure as the observed DC pairs (p<0.0009). This replication study using an independent cohort generated using different technology (microarray), indeed reinforces the robustness of our methodology and findings, and lends confidence to proceed with further downstream analyses.

### DC network contains region-exclusive interactions, and few hubs of AD dysregulation

For each inter-region BR1-BR2 comparison, we sought to identify whether the DC genes (**Table 1**) were exclusive to or shared among the two regions. We detected only 7%-21% gene overlap between the BR1 vs. BR2 DC genes (**Suppl. Fig. 3**), with the least vulnerable FP-STG brain region pair having the fewest gene overlaps of 7%, whereas the most vulnerable PHG-IFG having the highest 21% overlap (**Suppl. Fig 2**). Next, we wanted to check if these common genes in each inter-region analysis had the same gene neighbours in both brain regions, and found it not to be the case surprisingly (**Suppl. Table 2**).

Quantifying the pool of genes each brain region, say FP, uses to interact with three other brain regions under study (STG, PHG, and IFG), we realized that 10-50% of interactions are exclusive and only 20-30% interactions are shared across at least two regions (**Fig. 2a**). It seems a complex interplay between genes and region-specificity influences the activity of genes and their involvement in disease pathology. Together these analyses reinforce the importance of focussing on multiple brain regions and studying gene-gene interactions to understand AD aetiology.

**Figure 2.**
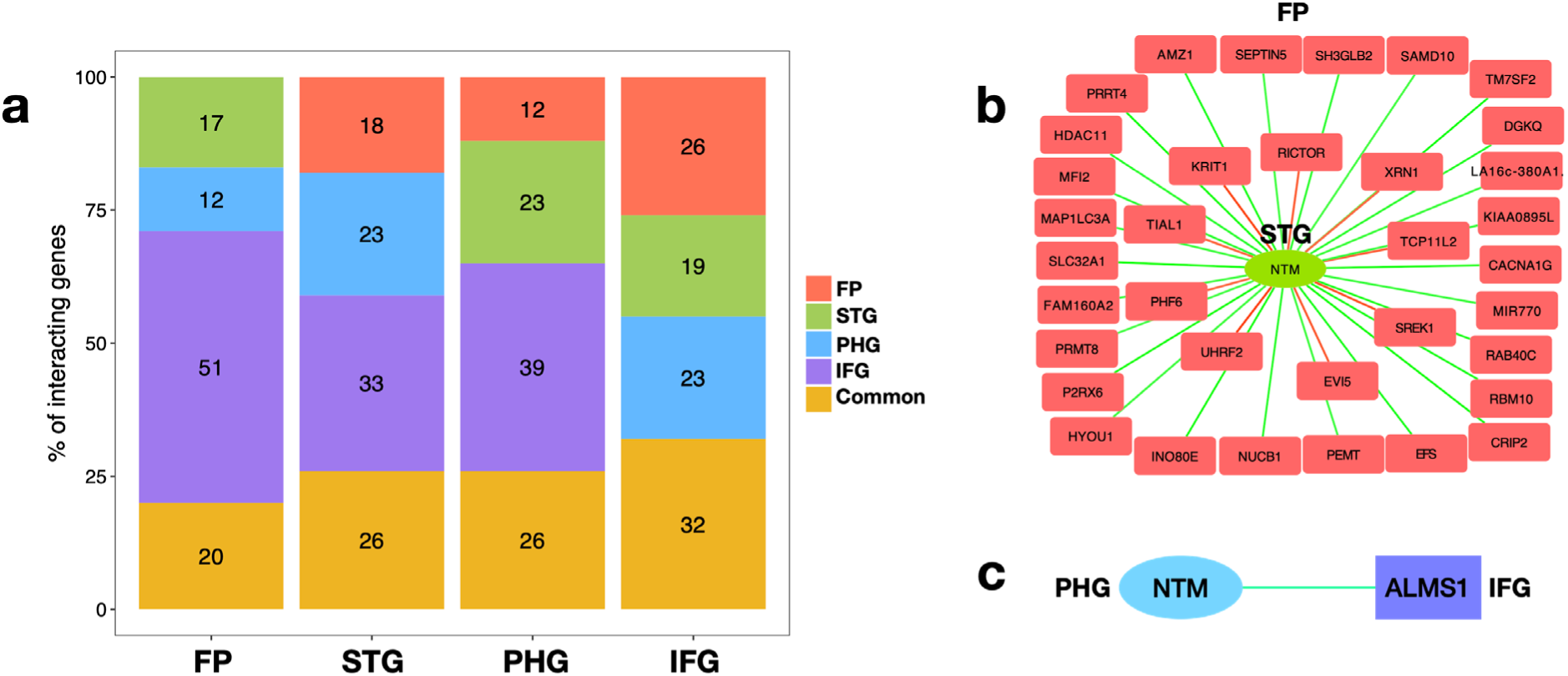
In the DC network, each brain region uses a unique gene profile to interact with other brain regions, and only a few genes act as hub genes. a. Stacked bar graph denotes that each brain region has an exclusive set of genes (12-51%) for interacting with other brain regions in the DC network. b. Hub gene *NTM* from STG is DC with 35 genes in FP. The edge colour represents a z-score. Green represents positively gained, and red represents negatively gained. c. In this network, *NTM* from PHG is differentially correlated to only one gene *ALMS1* in IFG.

After finding the exclusivity of inter-brain interaction, we aimed to identify the hub genes which participate in a large number of DC relations, and test if they are shared or exclusive across inter-region analyses. We are interested in gene hubs as they can underpin connectome hubs, which are well-connected brain network nodes that form a vital communication point for cohesive neuronal dynamics^10^. When examining the degree (number of DC interaction partners) of each gene for each inter-region comparison (**Suppl. Fig. 4**), we observed only a few hub genes, with more than 50% being non-hub having unit degree (single DC interaction). Moreover, the highest degree of hub gene in each brain region ranges from 20 (FP in FP-STG) to 113 (IFG in FP-IFG) (**Suppl. Table 3**). However, we noticed that a few hub genes with the highest degree are lncRNAs, pseudogenes, and antisense RNAs. Since their functionality is not well documented, we decided to select the protein-coding genes for our hub-gene analysis (**Table 3**). Interestingly, gene *ZKSCAN1* (Zinc Finger With KRAB And SCAN Domains) is found to act as a hub gene for different inter-region comparisons. *ZKSCAN1* is reported to have a role as a transcription factor that modulates GABA type-A receptor expression in the brain^26^. Exploring DC partners of ZKSCAN1, we realized it has region exclusive partners mostly. Two of its DC partner genes, SGK2 (Serum/Glucocorticoid Regulated Kinase 2) and TCF12 (Transcription Factor 12), are found in four inter-brain region comparisons (FP-IFG, STG-PHG, STG-IFG and PHG-IFG), where the DC edge ZKSCAN1-SGK2 is positively gained in all four and ZKSCAN1-TCF12 is negatively gained in all four analyses. While SGK2 is known to regulate ion channel transport and transport of glucose, metal ions^27^, etc., its involvement in AD is not known. On the other hand, TCF12, required for the initiation of neuronal differentiation^28^, is known to be dysregulated in AD^29^. Moreover, *NTM* (Neurotrimin), *LZTS1* (Leucine Zipper Tumor Suppressor 1), *FSD1* (Fibronectin Type III And SPRY Domain Containing 1), etc., are among a few other hub genes detected. The many DC partners of the hub gene *NTM* from FP-STG comparison is shown in **Fig. 2b**. Interestingly, in PHG-IFG comparison, *NTM* in PHG is DC with only one gene, *ALMS1* (*ALMS1* Centrosome And Basal Body Associated Protein) in IFG (**Fig. 2c**). This observation highlights that even the same gene from different regions have distinctive interactions. We reinstate that the same genes in multiple brain regions will have differing degrees of coupled expression with genes from another region, possibly due to the different spatial and molecular context they are in, and potentially have dominant region-exclusive effects.

**Table 3.**
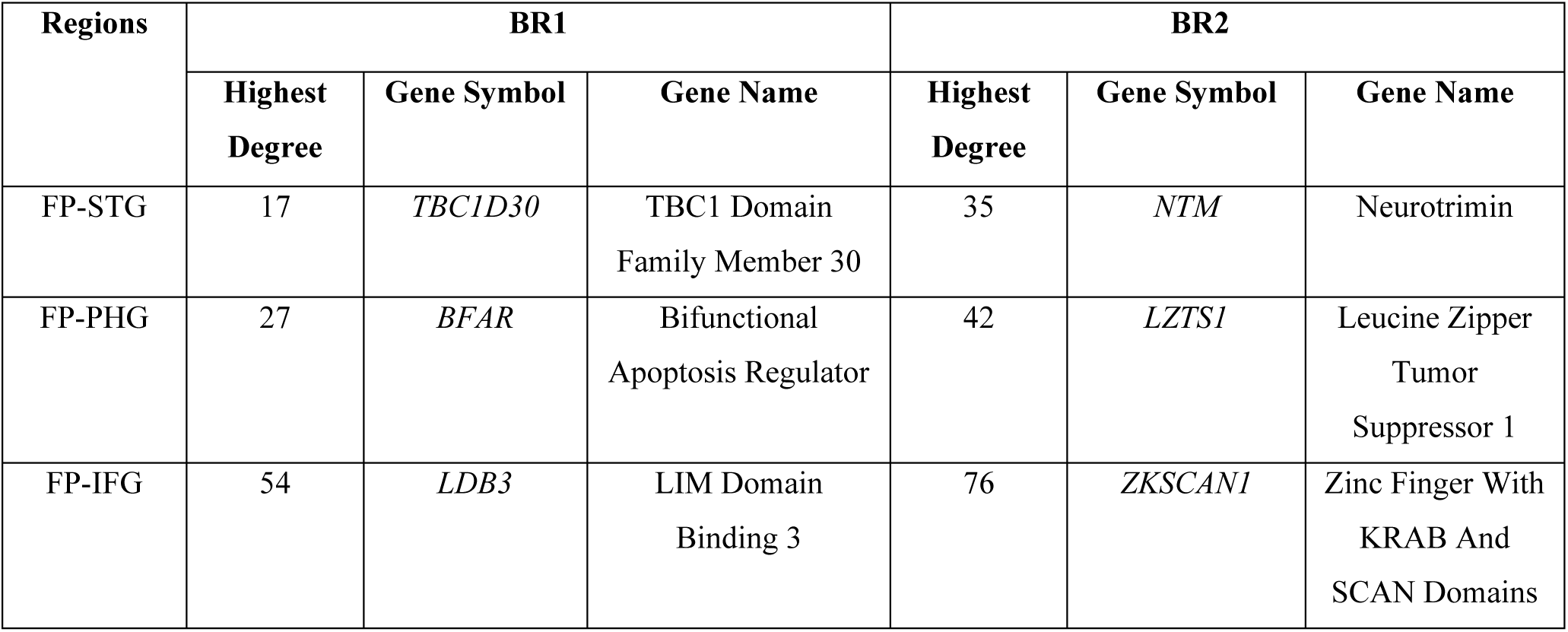
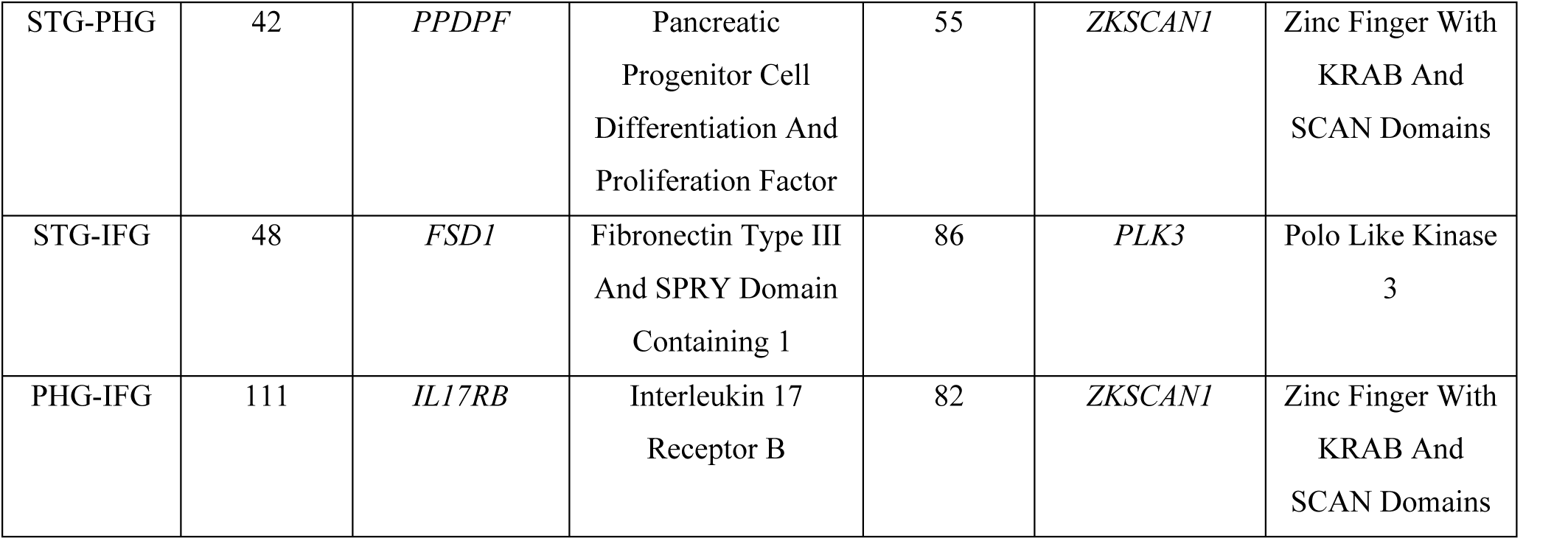
Hub protein-coding genes along with their degree from inter-region comparison (see also **Suppl. Table 3**)

### Bipartite network clustering reveals pathways with disrupted inter-region connectivity

DC genes are expected to provide valuable insights into the underlying biological processes of the clinical development of AD. To identify such biological processes, we partitioned the DC network into smaller bipartite (two-region) communities using the Louvain algorithm, such that genes within each community are more tightly connected among themselves than with genes in other communities. We excluded the “communities” comprised of singleton gene pairs, and finalized 19-34 communities per inter-region comparison, each of size of at least 20 genes. (**Fig. 3a, Suppl. Table 4**). Biological processes and pathways are controlled by vast interacting molecules whose expression levels are frequently co-regulated or co-expressed. After identifying tightly correlated DC modules, we performed over-representation analysis (ORA) to test if a set of DC genes is enriched for genes belonging to known Gene Ontology (GO) categories.

**Figure 3.**
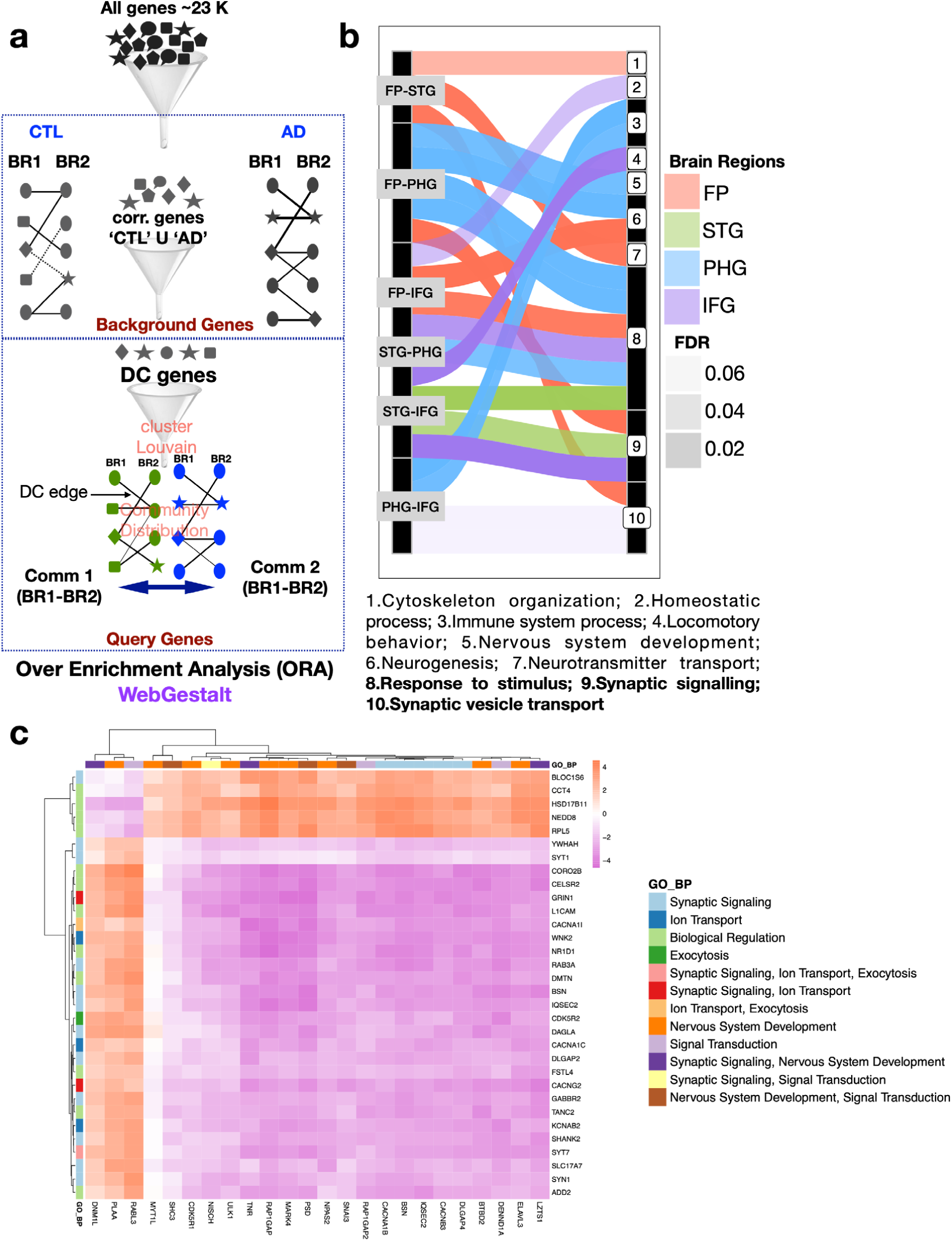
Bipartite DC modules provide insights into the inter-region biological processes affected by AD. a. Schematic diagram represents that correlated pairs from AD and CTL are used to calculate DC gene pairs, forming a bipartite network. This network is further partitioned into communities using the Louvain method (See Methods). Using the community genes as query genes and background gene list, over representation analysis is performed (see Methods). b. Alluvial plot represents the significant biological processes (BP) enriched in communities. The thickness of the edge represent the number of communities enriched for each BP for each brain inter-region comparison. c. Network connectivity between the enriched genes in com715 of FP-PHG is represented in the form of a heatmap, where the colour of the cell represents the z score of DC. Further, the GO-BP associated with each gene is represented using a colour label (GO-BP names are shortened, and their full names are in **Suppl. Fig. 6**).

Enrichment tests for GO biological processes (GO-BP) showed that most communities are enriched for response to stimulus, synaptic signaling, and transporter activities (**Fig. 3b**). The rest of the enriched functional profiles are highlighted in **Suppl. Fig. 5**. Among all the communities enriched, gene sets of community number 715 (hereafter referred to as com715) from both the brain regions in the FP-PHG pair were enriched for all three GO categories and KEGG pathways, with synaptic signaling being very prevalent. The top ten GO-BPs enriched from both brain regions are displayed as a hierarchical clustering tree along with the genes enriched for each process (**Suppl. Fig. 6**). Interestingly, GO-BP ‘Synaptic signaling’ and its related terms have different enriched gene sets from FP and PHG, respectively, except for five genes (*BSN, CACNA1B, GRIN1, IQSEC2*, and *SYNGAP1*). This reinforces our previous observation that every brain region’s dysregulated gene sets are distinctive. Among these five genes, *BSN* (Bassoon) is a component of the presynaptic active zone (AZ) involved in organizing the presynaptic cytoskeleton^30^. In contrast, voltage-dependent N-type calcium channel subunit alpha-1B (*CACNA1B*) mediates the ingress of calcium ions (Ca2^+^) into excitable cells, thus controlling the neurotransmitter release from the presynaptic compartment^31^. On the other hand, *GRIN1* encoding the essential subunit GluN1 that is present in all NMDARs (N-methyl-d-aspartate, receptors) found in the postsynaptic membrane, regulates the flow of Ca2^+^ through the channel^32^. Lastly, *IQSEC2* (IQ Motif And Sec7 Domain ArfGEF 2) and *SYNGAP1* (Synaptic Ras GTPase Activating Protein 1) form components of the postsynaptic density at excitatory synapses and are critical for the development of cognition and proper synapse function^33,34^. Except for *GRIN1*, none of the other genes are till date reported to be involved in AD pathology^35^. Since all these genes are involved in maintaining synaptic signaling, dysregulation of these genes will lead to neurodegeneration in a variety of disorders, including Alzheimer’s disease. As from our result, it is clear that these genes are differentially correlated in AD; it will be interesting to study whether the gene and protein expression levels of BSN, *CACNA1B, IQSEC2*, and *SYNGAP1* are altered in AD and, if so, how they contribute to AD pathology.

Further, we depicted the DC relation between only the synaptic signaling annotated genes in com715 in **Fig. 3c**. *WNK2* (WNK lysine deficient protein kinase 2) from FP and *LZTS1* (Leucine Zipper Tumor Suppressor 1) from PHG have the highest degree in PHG and FP, respectively. As evident from ‘The Human Protein Atlas’^36^, both genes and their corresponding proteins are expressed in the cerebral cortex. Studies have shown *WNK2* is present in cerebral cortex as well as cerebellum of mouse brains, enriched in neurons, and a regulator of GABAergic signaling^37^. Recently, it has been reported that *Lzts1* is associated with microtubule formation, contributes to the increasing intricacy of the cerebral architecture during evolution in mouse, and is mainly enriched in glial cells^38^. Their links to the brain is a motivation to find their connection with AD pathology. Moreover, we noted that most of the gene-gene correlations are lost in AD compared to CTL in com715. This probably indicates that synaptic signaling between FP and PHG is disrupted in AD.

### Genetic factors underpinning inter-brain region AD communication

Our DC analysis framework can help recapitulate current genetic/molecular understanding of AD pathology and open up new avenues to enhance this understanding further. Over the last decade, Genome-wide Association Studies (GWAS) have revealed many risk loci for AD, implicating many potential causative genes and SNPs (single nucleotide polymorphisms), beyond the well-established APOE association^39,40^. After identifying DC genes and the gene sets from the bipartite community, we wanted to check if these genes or gene sets are enriched in AD-GWAS signal using MAGMA, a tool for analysing GWAS data (**Suppl. Fig. 7**). Gene analysis yielded 208 significant AD-GWAS signal enriched genes, i.e., genes for which SNPs in their genomic vicinity are enriched for AD GWAS associations. A small set of genes overlap between 208 significant AD-GWAS signal enriched genes and DC genes from all six brain inter-region comparisons. However, interestingly our previously identified hub-gene *ZKSCAN1* is also enriched for AD-GWAS signal, making it an excellent novel candidate for AD pathogenesis that can be studied further in the context of its DC partners. Other than that, *CARF* (Calcium Responsive Transcription Factor) and *PLEKHA1* (Pleckstrin Homology Domain Containing A1) also enriched for AD-GWAS signals; being involved in the rewiring of gene coexpression networks in AD and further based on their functional relevance in the brain, they may also be considered as promising candidates for AD pathogenesis.

We then performed gene set analysis using MAGMA on 302 gene sets belonging to 151 communities, and two gene sets, namely com349 (present in FP of FP-IFG comparison) and com869 (present in IFG of FP-IFG comparison), were found to be significant. com869 comprises 54 genes, of which genes *NDUFS2, RAPGEF2*, and *PLEKHA1* are found to be significantly enriched for AD GWAS signal. Performing ORA on com349 gene set we found it to be further enriched at FDR 5% for nuclear factor (NF)-kappa β signaling pathway using KEGG as functional database. NF-κβ signaling pathway helps balance between learning and memory via its effect on synaptic plasticity, and disruption of this signaling module leads to neuroinflammation, oxidative stress, microglial activation, and apoptosis, all of which could promote AD^41,42^. Further, using the 208 significant AD-GWAS genes as functional database category and performing ORA revealed that six communities, four from FP and IFG interaction, and two from STG and PHG interactions, are enriched for AD-GWAS signals **(Fig. 4a).**

**Figure 4.**
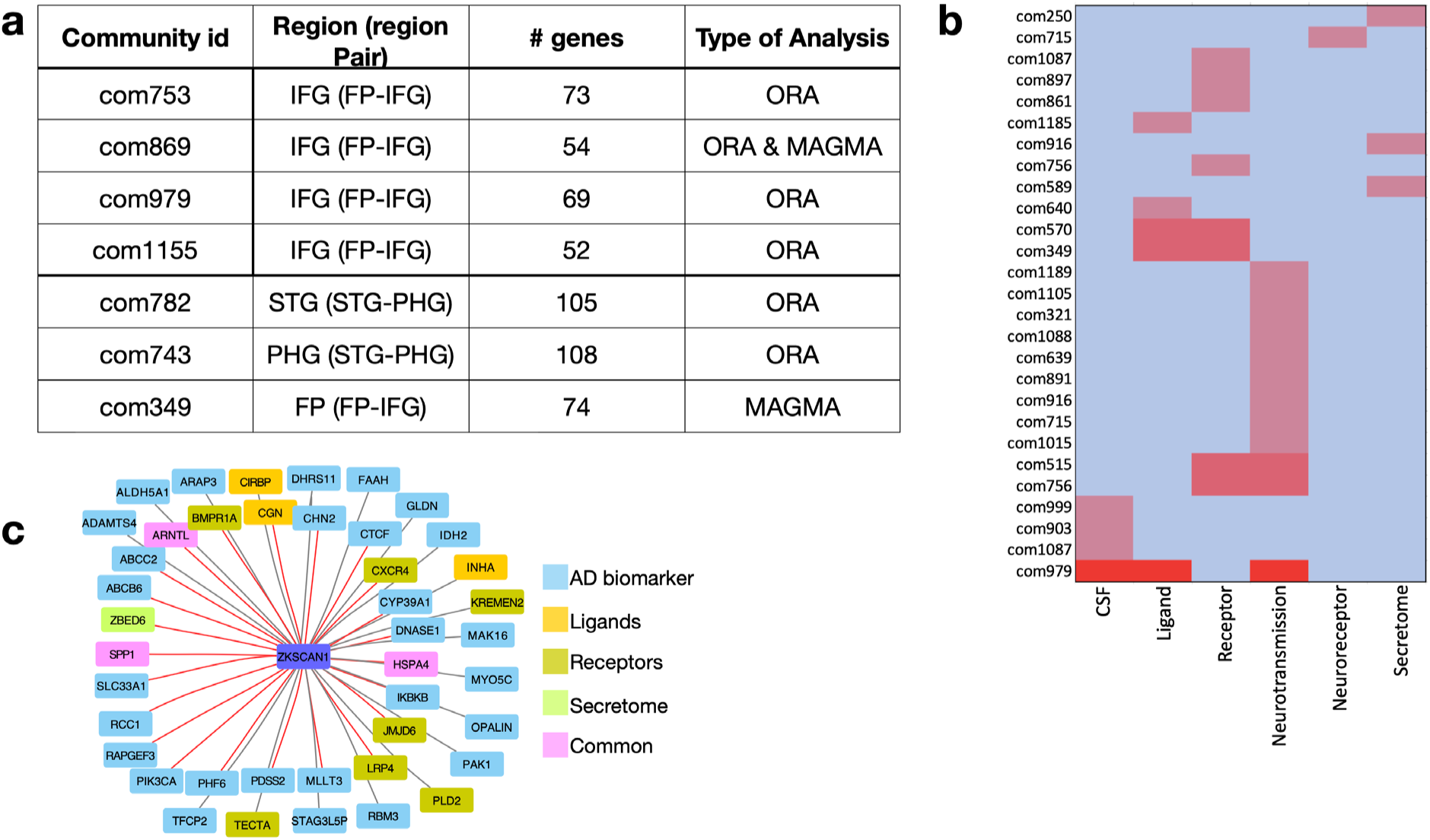
Genetic factors and signaling molecules underpinning brain inter-region communication and disease dysregulation. a. Communities (gene sets) enriched in MAGMA and ORA for genetic factors associated with AD. b. Heatmap indicating the type of functional categories for which a gene set is enriched. Gene sets (indicated with community number, details in **Suppl. Table 5**) are represented in the Y-axis. c. DC partners of *ZKSCAN1* are composed of AD biomarkers and signaling molecules. (edges in gray scale represents positively gained edge and in red scale represents negatively gained edge, according to z score)

### Distribution of signaling molecules in brain inter-region communities leads to molecular hypothesis of AD dysregulation

We predicted the rewiring of the gene network in AD via de-coupling and re-coupling of genes across different brain regions using RNA-sequencing gene expression data. However, the molecular mechanisms supporting this functional organization and re-organization remain elusive. Here, we tried to predict the functional nature of genes, which act as nodes of the rewired gene network. We selected those signaling molecules which are essential for communication between cells or regions near or far, such as ligand-receptor molecules, CSF markers, secreted proteins (secretome), and neurotransmitters-neuroreceptors (neurotransmission), and checked whether they are over-represented in communities derived from bipartite network partition. Using the above-mentioned molecules as functional categories in ORA, we could uncover 27 biologically meaningful, significant gene sets belonging to 23 communities at 5% FDR (**Fig. 4b**).

Gene sets from FP-IFG are most enriched for molecules responsible for communication. This may be due to their close proximity as both FP and IFG are located in the frontal cortex. We also found enrichment of secretome (com916), CSF (com979, com1087), ligand (com979) and receptor (com756, com979, com1087) molecules. Across all DC communities, 35 ligands and 61 receptors are found along with only 7 CSF and 9 secreted proteins. Finally, a total of 73 neurotransmitter genes out of 466 known neurotransmitters are found in different gene sets; and it is quite interesting that 12 gene sets while all being enriched for the same neurotransmission set of genes, do not overlap significantly in the specific neurotransmitters they contain, thereby reinforcing the exclusivity of the DC patterns to different regions and modules. Brain cells are known to communicate by passing neurotransmitters at the synapse, thereby transferring information throughout the brain. Noteworthy is that the previously determined com715, which is enriched for synaptic signaling and ion transport, is also enriched for neurotransmission in the FP side and neuroreceptor on the PHG side. The specific signaling molecules and interactions within the DC module com715 can thereby lead to specific hypothesis of how this module is affected in multiple regions in AD

## Discussion

We have presented here a new DC-based approach to study gene-gene correlation across brain regions and find the gene pairs that are functionally coupled and contribute to AD pathology. These analyses based on brain inter-region enable us to find the genic effect of one brain region on another brain region. We could further decipher genes that are not yet designated as AD biomarkers, but from our analyses, we could clearly observe that they are involved in gene pair rewiring in AD compared to CTL. Partitioning DC network into robust communities highlighted that gene pair rewiring is tightly linked to synaptic signaling and ion transport. Enrichment of these communities further for AD-GWAS signal as well as signaling molecules, helped us to build mechanistic hypotheses supporting the brain molecular connectivity. Further, hub gene analysis resulted in unveiling *ZKSCAN1* as key DC gene for most of the brain inter-region comparisons.

While this approach presents new facts about brain region functioning, particularly in AD pathology, it is worth pointing out that the results are limited by the brain regions for which data is available and further should be viewed as *in-silico* driven hypotheses that require experimental validation. We will get a better view once genomic data specific for brain regions is available, and experiments are pursued in future around the most promising lead genes/pathways from this study. Further, while CTC helps to reduce confounding effects of cellular composition, the CT frequencies are estimated and that too only for four major brain cell types. Nevertheless, we get meaningful results that are not enriched for CT specific marker genes. Despite some caveats, the result showing gene-pair rewiring across brain inter-region is of much interest and may help to study AD pathology in a new light.

From our analysis we realized that more than one aspect of synaptic function are affected such as synaptic vesicle trafficking, trans-synaptic signaling, chemical synaptic transmission, and regulation of synaptic plasticity. Interestingly, we noted that even if two brain regions e.g., FP and PHG, are coupled for synaptic signaling, they use a different set of genes to exert their effect. Downregulation of synaptic genes in AD and their specificity to brain regions have been previously reported^43^. However, the usage of different synaptic genes by different brain regions and their functional coupling across brain regions have not been reported before. Shifts in harmony of brain molecular connectivity involving synaptic genes is expected to compromise communication processes between brain regions leading to neurodegeneration, thus causing Alzheimer’s disease.

There remains a lack of detailed mechanistic knowledge about how the brain neural network is controlled. This is complicated by the fact that different brain regions are affected variedly by AD pathology, adding a spatial element to the disease. However, using the DC framework, we could identify communities of genes, whose gene network architecture between brain regions is altered in AD group relative to controls. We tried to hypothesize and visualize the cause and consequence of gene network dysregulation in AD pathogenesis using these communities of genes. We used SNP-based enrichment analysis and customized gene set analysis to confect our hypothesis. We found that 33 communities are enriched for several molecular factors such as AD biomarkers, CSF, neurotransmitters, ligand-receptors, AD enriched SNPs, and secretome proteins. Interestingly, com715 is enriched for neurotransmitters on the FP side, whereas on the PHG side, it is also enriched for neurotransmitters but not at a significant cut-off. This reinforces our finding that genes in com715 are enriched in the synaptic signaling biological process. Out of 35 communities, 12 communities are enriched for neurotransmitters. This probably indicates that synaptic signaling is most compromised during genetic rewiring. All these analyses help us to hypothesize possible mechanisms around inter-region regulation. It seems that a milieu of regulatory elements maintains the gene network of the brain underlying the neuronal network, and disruption of any or all factors rewires this network in AD patients.

Finally, hub gene analysis and AD SNP enrichments revealed that *ZKSCAN1*, located in chromosome 7, is a prominent node in the gene network functionally connecting different brain regions and has 310 unique DC partners. *ZKSCAN1*, when located in IFG, differentially correlates with the largest number of genes in other regions in AD. The pairing of *ZKSCAN1* with genes from different regions is either positively or negatively gained in AD compared to CTL. Previous literature indicates that *ZKSCAN1* can act as a transcription factor^44^. Hence, we looked for a known *ZKSCAN1* motif in the transcription start site of genes pairing with *the ZKSCAN1* gene using the HOMER motif analysis algorithm. However, none of the genes participating in DC relation with *ZKSCAN1* has its’ corresponding transcription factor motif. This suggests that *ZKSCAN1* uses a different mechanism to maintain correlation with genes from different brain regions, or there is not sufficient statistical power to detect *ZKSCAN1* motifs in its DC partner genes. Conversely, many of the *ZKSCAN1* DC partners are AD biomarkers (34), ligand (5), receptors (7) and secreted proteins (2). This indicates a plausible way how *ZKSCAN1* by controlling these signaling molecules (**Fig. 4c**) enacts its’ role in AD pathogenesis.

Comprehending AD pathology is not easy, however understanding brain connectivity alterations can give a better perspective. While functional and structural brain connectomes with respect to AD have been studied for a while now, focus on the molecular basis of these connectomes (molecular connectivity) is rare. Our inter-region DC framework addresses this gap by enlightening us with new findings and hypotheses on how AD affects the coupling between genes or biological processes in different brain regions. These results demonstrate the value of brain inter-region analysis in AD, and encourage its application to different neurological diseases and extension to inter-organ/inter-tissue analysis to understand the molecular connectome of the whole body.

## Materials and Methods

### Data collection

The covariate-adjusted RNA-sequencing data with the following synapse ids - syn16795931 – Brodmann Area (BM10) – frontal pole (FP), syn16795934 - BM22 - superior temporal gyrus (STG), syn16795937 - BM36 - parahippocampal gyrus (PHG), syn16795940 – BM44 - inferior frontal gyrus (IFG), were downloaded from AD Knowledge Portal – The Mount Sinai/JJ Peters VA Medical Center Brain Bank cohort (MSBB) study^45^ (10.7303/syn3159438). The pre- processed data is corrected for library size differences using the trimmed mean of M-values normalization (TMM method – edge R package) and linearly corrected for sex, race, age, RIN (RNA Integrity Number), PMI (Post-Mortem Interval), sequencing batch, exonic rate and rRNA (ribosomal RNA) rate. As in the earlier study^45^, normalization was performed on the concatenated data from all four brain regions to avoid any artificial regional difference.

The clinical (MSBB_clinical.csv) and experimental metadata (MSBB_RNAseq_covariates_November2018Update.csv) files available on the portal are used to classify the samples into control (CTL) and Alzheimer’s disease (AD) based on CERAD score (Consortium to Establish a Registry for AD; funded by NIA, 1986)^46^. CERAD score 1 was used to define CTL samples, and 2 (‘Definite AD’) was used for defining AD samples. Probable AD (CERAD = 3) and Possible AD (CERAD = 4) samples were not considered for this study. Sample sizes are noted in **Suppl. Table 6**. These samples were divided according to the four brain regions (**Suppl. Table 7**). Further, we considered the brain regions two at a time for our analyses and selected the samples accordingly to handle missing data (**Suppl. Table 8**; note that not all individuals had all four brain regions sampled).

The genes in the gene expression data are denoted in the hg37 ENSEMBL gene identifier (ENS. ID) format. The initial analysis is performed using the ENS. ID. For the downstream analyses (visualization/enrichment), the ENS. IDs are mapped to the HGNC gene symbols using the comprehensive gene annotation file for Release 19 (GRCh37.p13) downloaded from Gencode - https://www.gencodegenes.org/human/release_19.html (h37).

The Wang et al., 2016 study^20^ ranked 19 brain regions for their vulnerability to AD based on how many genes in these regions are associated with disease status (DE genes) and disease traits like the accumulation of NFT and Aβ. The brain regions used in this study are sorted based on the same ranking, such as BM36: rank 1, BM44: rank 2, BM22: rank 7, and BM10: rank 14, with rank 1 being the most vulnerable region in AD, and other ranks being proportionately less vulnerable.

Data for replication in independent cohort is retrieved from HBTRC^25^. Two brain regions - visual cortex (VC, BM17) and dorsolateral prefrontal cortex (DLPFC, BM9) - comprising of 300 samples (116 CTL and 184 AD) was used for our study. Gene expression data has been linearly adjusted for these covariates: age, gender, RIN, Batch, PMI and pH. Missing value of any covariate has been imputed with the respective mean value. Adjusted data is subjected to CTC as is done for MSBB; and same protocol for DC analysis is followed in this case also (explained below).

### Cell Type Correction (CTC)

The expression of a gene in a bulk tissue can be captured by the proportion of different cell types in the tissue and the expression of the gene in these cell types. Our ideal aim is to remove the former contribution and study the latter to reveal cell-intrinsic changes in gene pair correlation structure between disease vs. control group. Towards this, we corrected the bulk gene expression data for cell-type proportions, which were in turn estimated from bulk data using a cellular deconvolution method. Specifically, we estimated the frequencies of four major brain cell types, astrocytes, microglia, neuron, and oligodendrocytes, using a cellular deconvolution method implemented in the getAllSPVs function from the CellCODE (Cell-type Computational Differential Estimation) R package^47^. CellCODE is a singular value decomposition (SVD) based reference-free method to perform cellular deconvolution. It only requires the RNA-seq expression matrix of a set of marker genes. Human marker genes (markers_df_human_brain data frame) for the four major cell-types were obtained from the BRETIGEA (BRain cEll Type specIfic Gene Expression Analysis) meta-analysis study^48^. CellCODE performs F-tests on the supplied set of marker gene expression data to identify robust marker genes i.e., marker genes, which are not differentially expressed between the disease group vs. control groups. Only these robust marker genes are used to estimate cell-type proportions. The cell-type corrected gene expression data is obtained by linearly adjusting the bulk RNA-seq data for the cell-type proportions estimated using CellCODE.

We evaluated two different methods of cellular deconvolution, namely BRETIGEA and CellCODE on the cortical gene expression data set from Patrick et al. 2020 to identify the best performing model, i.e., the model with the highest correlation with the Immunohistochemistry (IHC) estimated cell-type proportions – which can be considered the ground truth data. For BRETIGEA, we used the function call: brainCells(geneExpmatrix, nMarker = 20, species = “human”), where nMarker is the number of markers that will be considered for each cell type to build the model. For CellCODE, we used the getAllSPVs function with input arguments: data, dataTag, grp, method, and mix.par, to build the model. Data is the gene expression data of the marker genes, dataTag is a binary matrix (# marker genes (MG) X # cell-types) which indicates which marker genes are associated with which cell type, and grp is the CERAD classification of each sample considered. Mixed method at the CellCODE-suggested 0.3 mix.par cutoff was used. The models for BRETIGEA and CellCODE were built using different sets of top 20, 40, 80, 200, 500, and 1000 marker genes sets for each of the four cell types to arrive at the best model.

Through this analysis, CellCODE 80 MG (i.e, 20 MG each of the four major cell types) was identified as the best performing model and hence used for the cell-type correction. Using CellCODE, we built one cellular deconvolution model for each brain tissue. By correcting DC interactions for cell-type composition effects, the confounding influence of cell-type proportions on the differential correlation results is mitigated^49,50^. The effect of applying the cellular deconvolution model is projected in Suppl. Fig. 1a, where the change in the number of correlated edges before and after CTC is highlighted for both AD and CTL samples using a bar chart. CTC has reduced the number of gene pairs compared to before CTC.

### Differential Correlation (DC) analysis

We are interested in identifying gene pairs across brain regions whose correlation strength in the disease group (AD) is significantly different from that in the control group of individuals (CTL) and call such pairs as differentially correlated or co-expressed (DC) pairs.

Gene-gene spearman correlation coefficients (ρ or r) for each of the gene pair combinations possible across brain regions are calculated for the AD group and CTL group separately. The Spearman correlation p-values are corrected for multiple testing using the Benjamini-Hochberg (BH) FDR method, and the resulting BH corrected p-values are subject to a 1% FDR cutoff to identify statistically significant correlation coefficients. All gene pairs significantly correlated either in the AD or Control group are considered for the DC analysis^51^. Note that we are not considering gene-gene interaction within a particular tissue. The union of correlated gene pairs of AD and CTL groups for any inter-region comparison is referred to as *correlated pairs* throughout the manuscript. Only these correlated pairs are tested for DC.

We use the r.test function from the psych R package to test a gene pair for DC. The r.test function transforms the AD as well as CTL gene-gene correlation coefficient values obtained for each gene pair into their corresponding z scores, known as the Fisher’s r to z transformation. The difference between the Fisher z transformed correlation coefficients, divided by the standard error of the difference, yields the final z-scores and associated DC p-values to be tested. For any inter-region comparison BR1-BR2 (Brain Region 1-2), we subject the DC p-values of all correlated gene pairs in BR1-BR2 to multiple testing correction using the Benjamini-Hochberg FDR method, and use 1% FDR cut-off to report significant DC pairs. For any given inter-region comparison, the DC Dsyregulation Index is the ratio of the number of significant DC gene pairs detected for that region pair to the number of all gene pairs tested for DC (i.e., all correlated pairs) for the same region pair. Note that the sign of a (DC) z-score indicates whether a particular DC gene pair’s correlation coefficient increased (positive z score) or decreased (negative z score) in the AD group relative to the CTL samples.

In addition, to check whether the sets of DC gene pairs in two inter-region comparisons are similar, the Jaccard similarity index, which is the ratio of the intersection of two sets to the size of their union, was calculated^52^.

### Differential Gene Expression Analysis

In this study, Differentially Expressed Genes (DEGs) were identified from CTC (cell type corrected) bulk RNA-seq data using a Wilcoxon rank-sum test for each of the four brain regions. DEGs identified at FDR cut-off 0.05, 0.1, and 0.2 were used to check whether the DC relation between each gene pair is driven by DEGs or not.

### Identification of bi-partite (two-region) modules or communities

The set of gene pairs identified as DC for two given brain regions (BR1 and BR2) can be viewed as a bipartite (two-layered) network of DC relations. We are interested in identifying a module or community comprising one set of genes in the first region (BR1) and another in the second region (BR2) that participates in many DC relations among themselves. We would also prefer that the communities be tightly-knit modules such that genes within a community are more likely to be related to one another than they are to the rest of the network. These preferences can be expressed as a modularity objective function. The bipartite network can be partitioned into a collection of modules or communities that maximize this modularity function using a heuristic method called the Louvain method^53^. The cluster_louvain function under the R package igraph was used for this purpose. Using the ‘ modularity ‘ function, we calculated the modularity score for each bipartite network (inter-region DC gene set). FP-STG has the highest modularity score of 0.95 and PHG-IFG the lowest, 0.63. To detect the communities enriched for significant Gene Ontology (GO) biological categories and pathways, we set a threshold that at least 20 genes must be present in each community. Partitioning each DC gene pair list from each inter-region comparison resulted in multiple communities. Each community comprises two gene sets, one from BR1 and one from BR2.

### Over Representation Analysis (ORA)

Over Representation Analysis (ORA) is a method that tests if genes from pre-defined functional sets (such as those belonging to a specific GO term or KEGG pathway) are enriched or over-represented (i.e., present more than would be expected by chance) in a given query set of genes.

To identify the potential biological functions associated with the gene sets or communities we identified, we performed ORA using the WebGestaltR package^54^. We performed this enrichment analysis only on communities of reasonable size (specifically those with at least 20 genes). A correlated gene list corresponding to each brain region (union of AD group and CTL group) was used as the background genes for this analysis, whereas the DC genes from each community acted as query genes. WebGestaltRBatch function was used to run the enrichment analysis so that the gene lists for multiple communities can be submitted at the same time. Under the ‘Functional database category’, Gene Ontology, GO (Biological process, cellular component, and molecular function), and pathways (KEGG & REACTOME) were selected for enrichment. We used FDR thresholds of 0.05 and used redundancy reduction methods (affinity propagation and weighted set cover) to find the most significantly enriched terms.

For the enriched communities, we ran the ORA with ShinyGO v0.66^55^ to generate the hierarchical clustering tree. This tree groups related GO terms together based on how many genes they share. The top 10 processes were selected for hierarchical clustering tree representation.

We also used customized functional categories, including genes enriched for AD GWAS signal, ligand-receptor molecules, CSF markers, secreted proteins, and neurotransmitters for ORA. The AD GWAS enriched genes are retrieved from MAGMA analysis (explained below). Ligand-Receptor pairs are assembled by combining the latest data of the year 2020 from GitHub repositories (https://github.com/LewisLabUCSD/Ligand-Receptor-Pairs). CSF markers are extracted from literature mining. Secreted proteins and neurotransmitters are downloaded using AmiGo^56^.

Further, for each hub gene from one brain region, we retrieved the connected genes from the respective brain region with which it interacted and performed over representation analysis (ORA) using WebGestalt, using the respective background gene list. However, none of the gene lists were enriched for any Gene Ontology (GO) processes. This prompted us to partition each bipartite network (inter-region comparison) into more meaningful gene communities.

### SNP enrichment analysis

GWAS studies have revealed numerous risk loci associated with AD, which harbor putative causative genes and variants. We aimed to check if DC genes or DC community gene sets are enriched for such GWAS-detected AD associations. The AD GWAS association signals in the form of SNP summary statistics are available for a comprehensive set of SNPs from a recent meta-analysis study of four major AD GWAS studies - the Psychiatric Genomics Consortium (PGC-ALZ), the International Genomics of Alzheimer’s Project (IGAP), the Alzheimer’s Disease Sequencing Project (ADSP), and UK Biobank (UKB). This study assessed the effect of 9,862,738 SNPs in 71,880 AD samples and 383,378 controls samples^12^. We would now like to test whether a given gene (or set of genes) is in the vicinity of many SNPs associated with AD in the above meta-analysis study. For this purpose, we use MAGMA, a tool for gene analysis and generalized gene-set analysis of GWAS data, in order to predict gene and gene-set level p-values using SNP-level p-values^57^. Inputs to MAGMA include SNP summary statistics of the meta-analysis study^12^ (downloaded from the CNCR/CTG LAB (Center for Neurogenomics and Cognitive Research/Complex Trait Genetics) website), and European 1000 Genomes reference data as described next. 22,665,064 SNPs retrieved from European 1,000 Genomes data files were first annotated to 19,354 genes from the hg19 genetic reference (human genome Build 37), using a 10 kb annotation window on either side of the gene. Next, using SNP p-value and European 1000 Genomes reference data, 18,445 genes were mapped to SNPs, of which genes significantly enriched for AD GWAS signal at FDR<0.05 (Benjamini Hochberg, BH p adjustment) were retained. Further, gene set analysis was performed on 302 gene sets, and significantly enriched gene sets for AD GWAS signal at FDR 0.05 (BH adjusted p<0.05) was reported.

## Code Availability

Code used in this study is provided here: https://github.com/BIRDSgroup/InterTissueDC.

## Acknowledgments

We thank members of our BIRDS (Bioinformatics and Integrative Data Science) research group, and IBSE (Center for Integrative Biology and Systems medicinE) for their feedback during presentations of this work. We thank Nitish Mahapatra, Himanshu Sinha, and Karthik Raman for their valuable inputs on this work. The research presented in this work was supported by Wellcome Trust/DBT grant IA/I/17/2/503323 awarded to MN.

