## Supplemental Information (Suppl. Tables/Figures) for "Alzheimer’s disease rewires gene coexpression networks coupling different brain regions"

#### Supplementary Tables

**Supplementary Table 1. Classes of DC pairs detected by our analysis.** For a DC gene pair, its class is based on the gene pair's correlation in the CTL and in the AD group (1 means significantly positively correlated, -1 means significantly negatively correlated, and 0 means neither).

| CTL | AD | Status/Class | z-score |
|---|---|---|---|
| 0 | 1 | Gained positive correlation | Positive |
| 0 | -1 | Gained negative correlation | Negative |
| 1 | 0 | Lost correlation | Negative |
| -1 | 0 | Lost correlation | Positive |

**Supplementary Table 2. Summary of the common genes shared by two regions in an inter-region comparison.** Gene neighbours of these common genes are also inspected for overlap. Identical gene pairs per comparison are also noted.

| Brain Region Pairs (BR1-BR2) | Total DC edges | BR1 DC genes | BR2 DC genes | Common Genes | Gene neighbours of common genes (DC edges) | | Common neighbour genes | Identical gene pairs |
|---|---|---|---|---|---|---|---|---|
| | | | | | BR1 | BR2 | | |
| FP-STG | 2961 | 2013 | 1844 | 260 | 354 (444) | 392 (472) | 26 | 1 |
| FP-PHG | 2629 | 1597 | 1409 | 261 | 387 (521) | 439 (606) | 46 | 14 |
| FP-IFG | 9962 | 4609 | 4235 | 1299 | 1795 (3392) | 1850 (3338) | 314 | 12 |

| | | | | | | | | |
|---|---|---|---|---|---|---|---|---|
| STG-PHG | 6274 | 2580 | 2549 | 754 | 1071 (2595) | 1229 (3023) | 356 | 18 |
| STG-IFG | 8179 | 3297 | 3548 | 968 | 1520 (3198) | 1409 (2932) | 287 | 14 |
| PHG-IFG | 12979 | 3642 | 4202 | 1348 | 2441 (7402) | 1780 (6279) | 624 | 31 |

*Note- BR1 and BR2 represents brain region 1 & 2 respectively for each inter-region comparison. For example, in FP-STG brain region pair, FP is BR1 and STG is BR2.This abbreviation is used in subsequent tables also. In the case of FP-STG, there are 260 genes common between 2 regions, which is 13% of 2013 (FP) and 14% of 1844 (STG) genes. These common genes have 472 connections (DC edges) from BR1 to BR2 and 444 from BR2 to BR1, i.e., 260 genes of BR1 interact with 392 genes in BR2. Among these gene neighbours, only 26 genes are common. Further, 1 gene which forms identical gene pair is basically a subset of 260 common & also 26 common neighbour genes.

**Supplementary Table 3. Hub genes along with their degree from inter-region comparison.** Degree of a gene refers to the number of its DC partners (for abbreviations, refer to the note above).

| Brain Region Pairs (BR1-BR2) | BR1 | | | BR2 | | |
|---|---|---|---|---|---|---|
| | Highest Degree | Gene Symbol | Gene Name | Highest Degree | Gene Symbol | Gene Name |
| FP-STG | 20 | RP11-418J17.1 | WARS2 antisense RNA 1 | 35 | NTM | Neurotrimin |
| FP-PHG | 41 | FAM86 B3P | Family With Sequence Similarity 86 Member B3, Pseudogene | 42 | LZTS1 | Leucine Zipper Tumor Suppressor 1 |
| FP-IFG | 61 | TTLL7-IT1 | lncRNA: TTLL7 Intronic Transcript 1 | 113 | CACYBPP1 | Calcyclin Binding Protein Pseudogene 1 |
| STG-PHG | 42 | PPDPF | Pancreatic Progenitor Cell Differentiation And | 81 | FAM86B3P | Family With Sequence Similarity 86 Member |

| | | | | | | |
|---|---|---|---|---|---|---|
| | | | Proliferation Factor | | | B3, Pseudogene |
| STG-IFG | 48 | FSD1 | Fibronectin Type III And SPRY Domain Containing 1 | 86 | PLK3 | Polo Like Kinase 3 |
| PHG-IFG | 111 | IL17RB | Interleukin 17 Receptor B | 82 | ZKSCAN1 | Zinc Finger With KRAB And SCAN Domains |

**Supplementary Table 4**. **Statistics of communities detected in each brain inter-region comparison.** Largest community size (number of genes) for each comparison is noted. The number of communities used for enrichment analysis, based on a minimum size of 20 genes, is also shown.

| **Brain Region Pairs** | **Total** | **Max Size** | **Modularity Scores** | **Community Size >= 20** |
|---|---|---|---|---|
| FP-STG | 1051 | 105 | 0.9543194 | 19 |
| FP-PHG | 737 | 225 | 0.8996255 | 19 |
| FP-IFG | 1235 | 651 | 0.8082181 | 34 |
| STG-PHG | 907 | 313 | 0.7288623 | 27 |
| STG-IFG | 1130 | 424 | 0.7629159 | 29 |
| PHG-IFG | 1057 | 766 | 0.6348189 | 23 |

**Supplementary Table 5. Explanation of gene set name abbreviation -** The nomenclature of a gene set name is defined by its region, brain inter-region pair (in BR1-BR2 format), and the corresponding community number. For example, IFG(FP-IFG)-903 refers to the IFG-side gene set of the DC community 903 detected in brain inter-region pair FP-IFG's analysis. The gene sets shown here are from **Fig. 4b of main text**.

| **Gene Set Name** | **Abbreviation** |
|---|---|
| FP(FP-STG)-1015 | com1015 |
| FP(FP-PHG)-250 | com250 |
| FP(FP-PHG)-715 | com715 |
| FP(FP-IFG)-1087 | com1087 |
| FP(FP-IFG)-1185 | com1185 |

| | |
|---|---|
| FP(FP-IFG)-1189 | com1189 |
| FP(FP-IFG)-515 | com515 |
| FP(FP-IFG)-570 | com570 |
| FP(FP-IFG)-639 | com639 |
| FP(FP-IFG)-756 | com756 |
| FP(FP-IFG)-861 | com861 |
| FP(FP-IFG)-891 | com891 |
| FP(FP-IFG)-897 | com897 |
| FP(FP-IFG)-916 | com916 |
| FP(FP-IFG)-979 | com979 |
| IFG(FP-IFG)-1087 | com1087 |
| IFG(FP-IFG)-349 | com349 |
| IFG(FP-IFG)-756 | com756 |
| IFG(FP-IFG)-903 | com903 |
| IFG(FP-IFG)-916 | com916 |
| IFG(FP-IFG)-999 | com999 |
| IFG(STG-IFG)-1088 | com1088 |
| IFG(STG-IFG)-321 | com321 |
| PHG(FP-PHG)-715 | com715 |
| PHG(PHG-IFG)-640 | com640 |
| STG(STG-PHG)-589 | com589 |
| STG(STG-IFG)-1105 | com1105 |

**Supplementary Table 6. Sample sizes (# of individuals) according to the CERAD category.**

| CERAD Score | CERAD Category | Number of individuals |
|---|---|---|
| 1 | Controls | 100 |
| 2 | Definite AD | 170 |

**Supplementary Table 7. Sample sizes (# of RNA-seq samples) in the four brain regions.**

| Brain Region | CTL | AD |
|---|---|---|
| FP | 84 | 146 |
| STG | 80 | 129 |
| PHG | 63 | 115 |
| IFG | 66 | 121 |

**Supplementary Table 8. Sample sizes (# of RNA-seq samples) used in the inter-brain region comparisons.**

| Brain Region Pairs | CTL | AD |
|---|---|---|
| FP-STG | 58 | 88 |
| FP-PHG | 59 | 77 |
| FP-IFG | 59 | 80 |
| STG-PHG | 56 | 74 |
| STG-IFG | 54 | 80 |
| PHG-IFG | 55 | 69 |

### Supplementary Figures

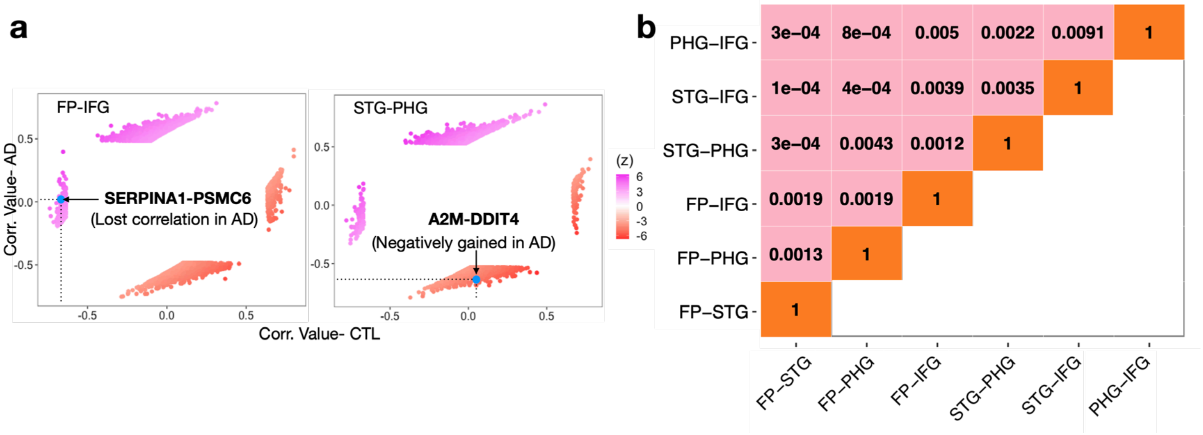


**Supplementary Figure 1. Summary of DC analysis across four brain regions.**

a. In two example inter-region comparisons, detected DC gene pairs form four distinct clusters in the scatter plot (see also **Suppl. Table 1**), representing different types of change between AD and CTL. Example DC gene pairs pointed in this figure illustrate two of these types/clusters of change, and that in **Fig. 1c of main text** illustrate the other two types.

b. DC gene sets across brain inter-region comparison share almost no similarity as indicated by Jaccard index.

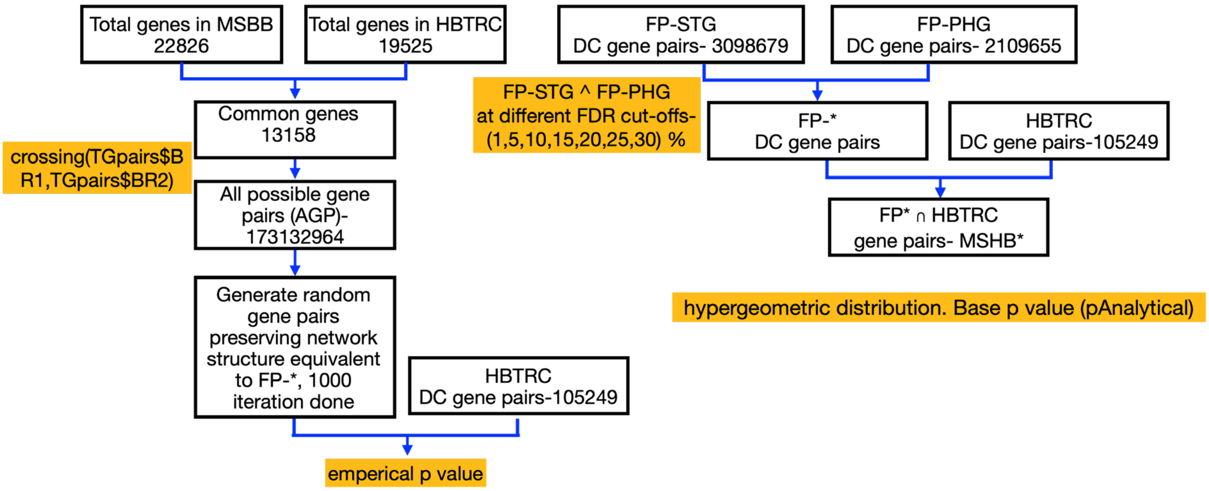


**Supplementary Figure 2. Schematic of replication testing of DC results found in our discovery cohort (MSBB) in an independent validation cohort (HBTRC).**

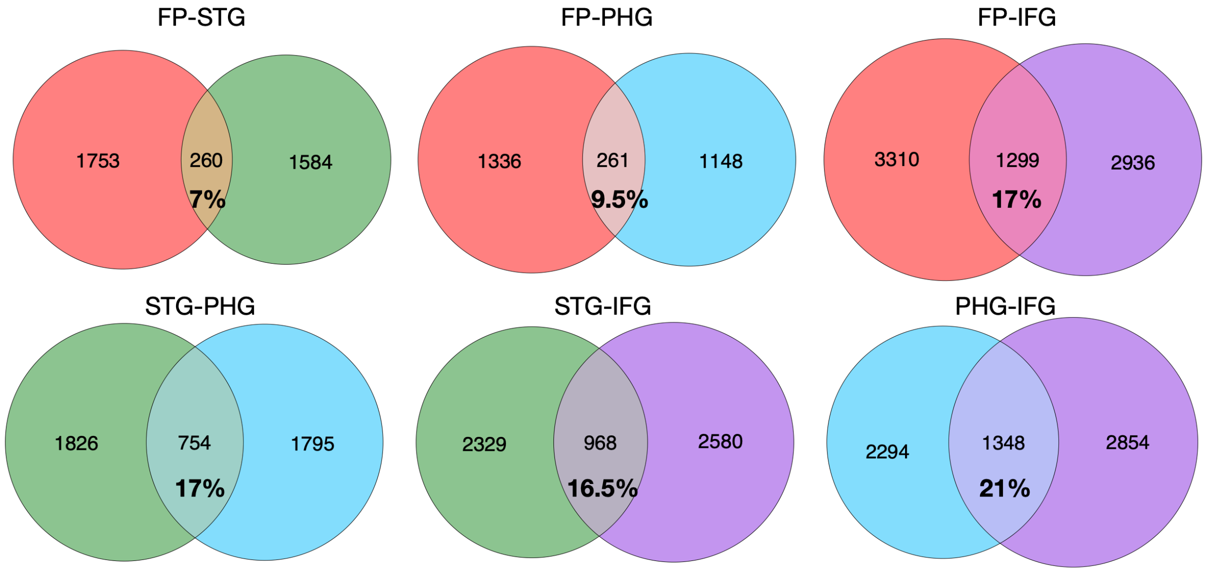


**Supplementary Figure 3. Number and percentage of common genes across each brain inter-region comparison.** See also **Suppl. Table 2.**

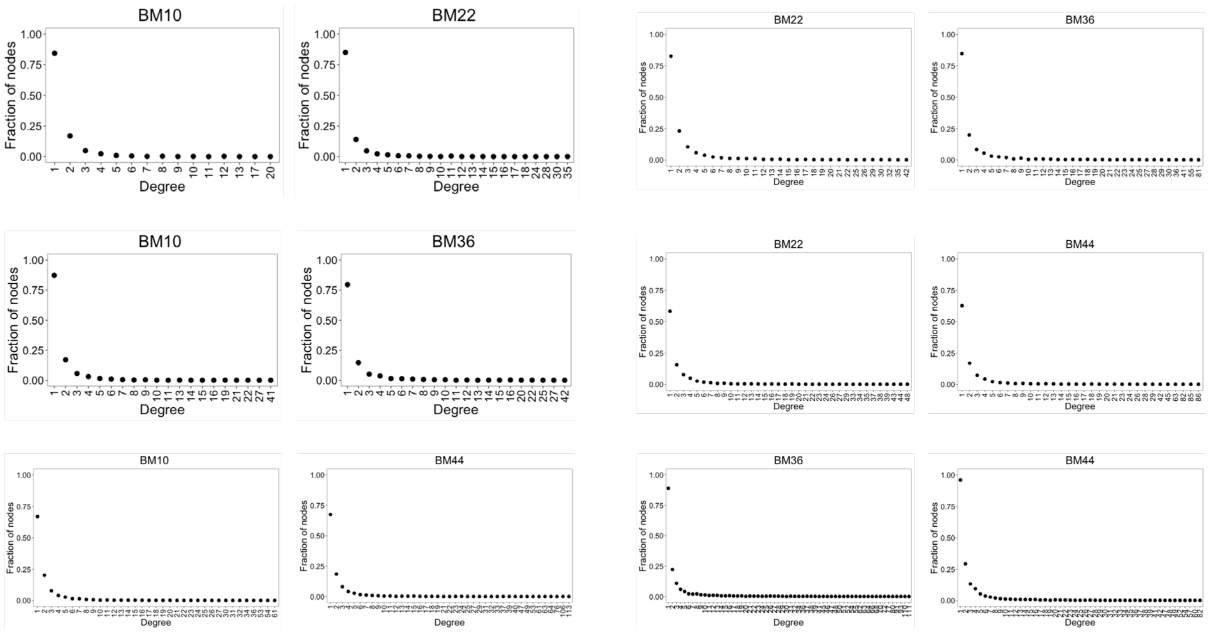


**Supplementary Figure 4. Degree distribution of genes in a brain region in each inter-region comparison.** A degree of a gene in an inter-region comparison is the number of DC relations/interactions that this gene participates in.

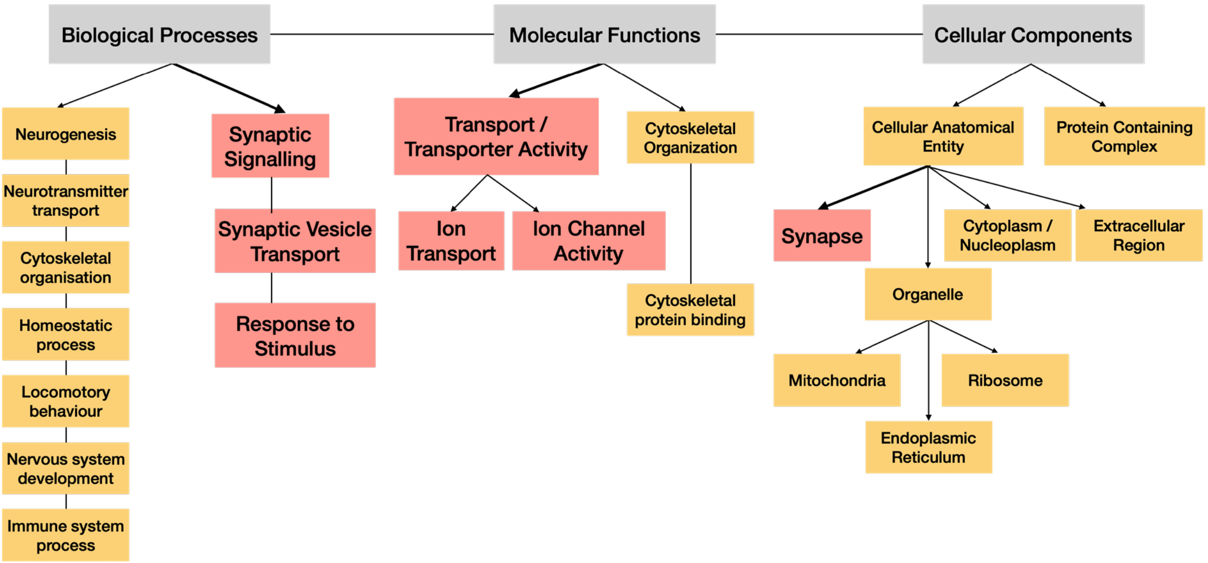


**Supplementary Figure 5. Summary of GO categories' terms enriched in our DC communities.** GO terms shown in a red box are enriched in most of the communities, whereas the rest are enriched in 1-2 communities.

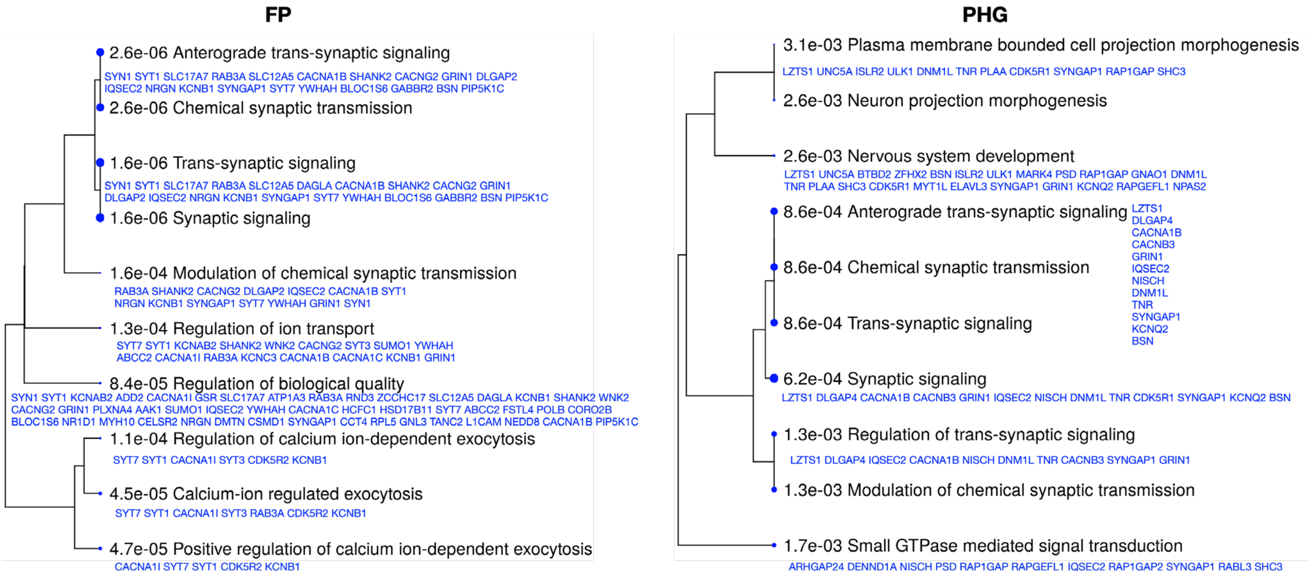


**Supplementary Figure 6. Summary of overlap among GO terms enriched in com715.** (left) GO terms enriched in the FP gene set of our DC community FP-PHG::com715 are shown, along with the com715 genes overlapping with the corresponding GO term. Similarity of these overlapping genes of different GO terms are used to build the hierarchical clustering tree shown. Size of a solid circle indicates the (adjusted pvalue based) significance of the corresponding enrichment. (right) Similar results for GO terms enriched in the PHG side of com715.

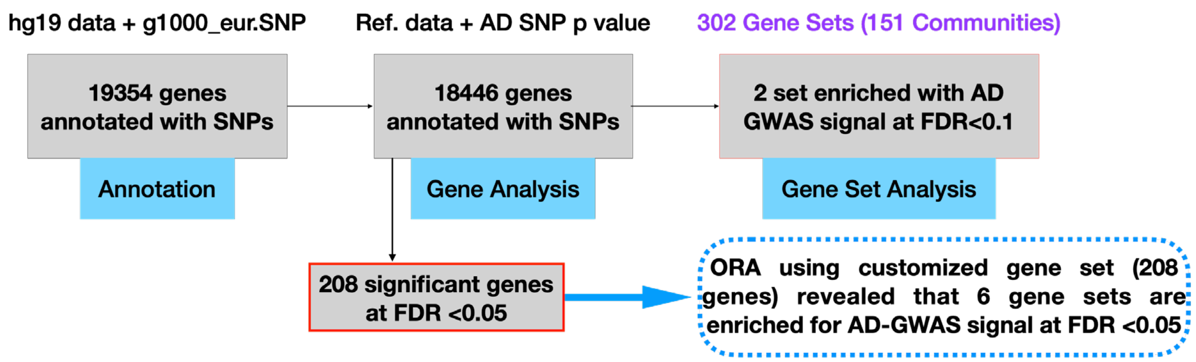


**Supplementary Figure 7. Enrichment of DC communities for GWAS signals - Workflow.** Schematic of the steps involved in MAGMA gene-level and gene-set-level analysis, and customized ORA (Over Representation Analysis) that tests whether gene sets are enriched for the 208 AD-GWAS-signal enriched genes (see also Methods). Each community yields two gene sets (from the two brain regions), which are then tested for enrichment.
